# A Network-Based Discovery of Prognostic Markers in Recurrent IDH wild-type Gliomas

**DOI:** 10.1101/2025.07.17.665414

**Authors:** Yang Liu, Jason Huse, Kasthuri Kannan

**Author notes:** Contributing authors.

## Abstract

**Background:** Isocitrate dehydrogenase wild-type (IDH wild-type) gliomas represents the most aggressive subtype of diffuse gliomas, characterized by therapeutic resistance and dismal prognosis. Despite advances in molecular classification, reliable prognostic biomarkers for these tumors remain limited, particularly for recurrent disease. This study aims to identify gene expression signatures associated with survival outcomes in recurrent IDH wild-type gliomas, with the goal of improving patient stratification and potential therapeutic targeting.

**Methods:** We analyzed gene expression data from 180 recurrent IDH wild-type glioma samples from the Glioma Longitudinal AnalySiS (GLASS) Consortium. Using multiple computational approaches including a novel network-based method (netSurvival) and various survival analysis techniques, we identified genes associated with patient survival outcomes.

**Results:** Our comprehensive analysis identify several gene expression markers that are significantly associated with survival outcomes in recurrent IDH wildtype gliomas. The AFT log normal model revealed that FN1, HIF3A, and EIF4B are associated with poorer survival (hazard ratios of 1.40, 1.49, and 1.54, respectively; p ¡ 0.05), while PTK2, CCND2, RAD51L3 RFFL, and MAX demonstrated protective effects (hazard ratios of 0.76, 0.78, 0.79, and 0.79, respectively; p ¡ 0.05). Five genes (KIF5C, LINC00632, B4GALNT3, HIF3A, and RAD51L3 RFFL) show significant differential expression between primary and recurrent tumors, with four having established functional roles in glioma pathobiology.

**Conclusion:** This study identifies a panel of gene expression markers with significant prognostic value in recurrent IDH wild-type gliomas. The differential impacts of these genes on survival outcomes provide insights into the biological heterogeneity underlying clinical behavior in these aggressive tumors. Particularly significant are the biomarkers associated with both survival outcomes and recurrence patterns, which may represent key drivers of disease progression. These findings represent an important step toward improved prognostic stratification and therapeutic targeting in IDH wild-type gliomas, addressing a critical unmet need in neuro-oncology.

## 1 Introduction

IDH wild-type gliomas constitute a distinct molecular entity characterized by aggressive clinical behavior and poor prognosis. Representing over 90% of glioblastomas (GBM), the most common and malignant primary brain tumor in adults, these tumors present a formidable challenge in neuro-oncology. Despite advances in multimodal treatment approaches combining surgical resection, radiotherapy, and chemotherapy, the prognosis for patients with IDH wild-type gliomas remains dismal, with a median survival of merely 12-15 months [1]. This poor survival rate highlights the urgent clinical need for improved stratification methods and novel therapeutic approaches for this devastating disease.

The molecular landscape of gliomas has been substantially refined in recent years, leading to the 2016 World Health Organization (WHO) classification integration of molecular parameters into the diagnostic algorithm. The identification of IDH mutation status as a fundamental molecular classifier has divided diffuse gliomas into distinct biological and clinical entities. While IDH-mutant gliomas generally exhibit more favorable outcomes, IDH wild-type gliomas demonstrate aggressive behavior and therapeutic resistance [2, 3].

The molecular heterogeneity of IDH wild-type gliomas presents both a challenge and an opportunity for biomarker discovery. Comprehensive genomic analyses have revealed several key molecular alterations that characterize these tumors. EGFR amplification occurs in 35-45% of IDH wild-type GBM, while CDKN2A/B homozygous deletion is observed in approximately 50-60% of cases [4]. TERT promoter mutations are present in over 80% of IDH wild-type GBM, serving as a key diagnostic marker that distinguishes them from lower-grade gliomas [5]. Currently, MGMT promoter methylation status remains the most clinically relevant predictive biomarker, with approximately 40% of IDH wild-type GBM exhibiting methylation that correlates with improved response to temozolomide treatment [6, 7].

Beyond genomic alterations, transcriptomic studies have provided valuable insights into the tumor microenvironment and cellular heterogeneity of IDH wild-type gliomas. Neftel et al. identified distinct cellular states and plasticity in glioblastoma that influence tumor progression and therapeutic resistance [8]. Ran et al. characterized the patterns of immune cell infiltration and their association with survival, highlighting potential immunotherapeutic targets [9]. However, these established biomarkers provide insufficient granularity for accurate prognostication and personalized treatment planning, underscoring a critical gap in our ability to identify targeted therapeutic approaches.

Despite these advances, current approaches to biomarker identification have often focused on isolated genetic alterations or protein expression levels without integrating comprehensive transcriptomic signatures with robust survival analyses. Advanced genomic and transcriptomic profiling has generated vast amounts of molecular data. However, a methodological gap exists in effectively translating this information into clinically actionable biomarkers that reliably correlate with survival outcomes.

In this study, we integrate gene expression profiles from recurrent IDH wild-type gliomas with patient survival data to identify potential prognostic biomarkers. This focus on recurrent tumors is particularly significant, as these represent the most challenging clinical scenario with even more limited treatment options and worse outcomes compared to newly diagnosed cases [10, 11]. Recurrent IDH wild-type gliomas often demonstrate heightened therapeutic resistance and altered molecular profiles compared to primary tumors, potentially revealing distinct survival-associated biomarkers [12, 13]. By employing network-based survival (netSurvival) approach and regularized Cox models, we aim to discover gene expression signatures that are critical for patient survival. Furthermore, we validate these findings through prognostic abilities and survival differences of the stratified groups via prognostic indexes (PIs).

The identification of reliable prognostic biomarkers for IDH wild-type gliomas holds significant promise for advancing precision medicine in neuro-oncology. Such biomarkers enable more accurate patient stratification for clinical trials, guide treatment decisions, reveal underlying biological mechanisms of tumor progression, and potentially uncover novel therapeutic targets. Ultimately, this research seeks to address the fundamental clinical challenge of improving outcomes for patients with one of the most aggressive and treatment-resistant malignancies in oncology.

## 2 Methods

### 2.1 Dataset and preprocessing

This study uses glioma data from the GLASS consortium. After preprocessing, gene expression TPM values and corresponding clinical outcomes are obtained for 180 IDH wild-type recurrent samples from 158 patients. Approximately 11.1% of the samples are right-censored.

Genes with more than 80% zero expression values and samples with missing data are removed. A five-fold cross-validation is performed to identify markers associated with survival. For each fold, the training and testing sets are normalized separately using log_2_(x + 1) transformation followed by z-score normalization. Due to the high dimensionality of the dataset (over 20,000 genes), a univariate log-rank test is applied to select the top 1,000 survival-related genes for subsequent analysis. Expression levels are dichotomized into high and low groups based on the median values.

### 2.2 Network construction and random walk for feature selection

As previously described [14], we apply one-dimensional hierarchical clustering to group samples for each gene based on their normalized expression values, identifying clusters of patients with the top 10% highest absolute log_2_ expression. These clusters form the nodes of a gene-gene interaction network, where edges are defined by shared patients between nodes, enabling the model to focus exclusively on gene-level interactions. To minimize the influence of indirect or complex relationships within the same gene, we randomly select one sample cluster per gene during each graph sampling iteration and repeat this process 1,000 times. We then apply a random walk algorithm to identify paths of connected nodes whose associated patient groups exhibit significantly different survival outcomes, as determined by a log-rank test comparing patients within the path to all others. Finally, we use Fisher’s exact test to identify prognostic markers, testing the null hypothesis that the presence of specific markers is independent of the significant survival-associated paths.

### 2.3 Regularized Cox Models

Due to the censored properties of time-to-event outcomes, such as overall median survival time, conventional methods like ANOVA are not appropriate for analyzing survival data in IDH wild-type gliomas. To address this, we employ three regularized Cox proportional hazards (PH) models, specifically Ridge Cox, Lasso Cox, and Elastic Net Cox, for feature selection to identify prognostic biomarkers in IDH wild-type gliomas. These models are well-suited for handling high-dimensional genomic data and account for censoring by modeling the hazard function over time.

The Ridge Cox model applies L2 regularization, which shrinks regression coefficients toward zero to reduce overfitting while retaining all features, making it effective for correlated predictors [15, 16]. The objective function for Ridge Cox regression is given by:

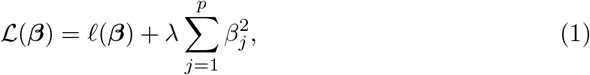

where *ℓ*(***β***) is the Cox partial log-likelihood, ***β*** = (*β*_1_, …, *β*_*p*_)^⊤^ are the regression coefficients, and *λ >* 0 is the regularization parameter.

In contrast, the Lasso Cox model uses L1 regularization to perform both shrinkage and feature selection by setting some coefficients to exactly zero, thus identifying a sparse set of predictors [15, 17]. The Lasso Cox objective function is:

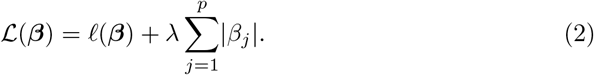

The Elastic Net Cox model combines L1 and L2 regularization, balancing the strengths of Ridge and Lasso to handle correlated features while selecting a subset of relevant biomarkers [18, 19]. The Elastic Net Cox objective function is:

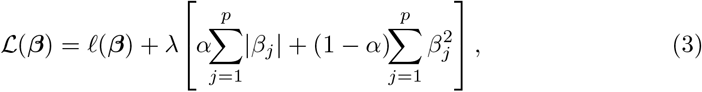

where *α* controls the balance between L1 and L2 penalties.

### 2.4 Machine learning models for survival outcomes

To evaluate the selected prognostic biomarkers for IDH wild-type gliomas, we employ several specialized survival analysis methods tailored to handle right-censored survival data.

The Cox PH model assumes a linear relationship between predictors and the logarithm of the hazard function, providing a robust framework for assessing the impact of selected biomarkers on survival outcomes [20]. The hazard function is modeled as:

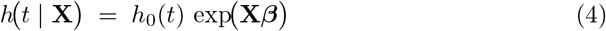

Where *h*_0_(*t*) is the baseline hazard function, **X** represents the covariates, and *β* are the estimated coefficients. As mentioned, the Elastic Net Cox model extends the Cox PH model by incorporating L1 and L2 regularization.

Random Survival Forests (RSF), an extension of traditional random forests, construct an ensemble of survival trees using bootstrap samples, effectively capturing complex, non-linear relationships between predictors and survival outcomes while accommodating censoring [21, 22]. RSF predicts the cumulative hazard function based on ensemble averaging of Nelson-Aalen estimators from individual trees.

Additionally, Support Vector Machines (SVM) adapted for survival data extend the standard SVM framework by integrating censoring information into the optimization process, modeling the relationship between high-dimensional predictors and time-to-event outcomes [23, 24]. The survival SVM formulation focuses on maintaining the correct ranking of survival times among pairs of patients.

To assess model performance, we used two primary metrics:

#### Concordance Index (C-index)

The C-index measures the model’s discriminative ability by calculating the proportion of all possible pairs of patients where the patient with the higher predicted risk experiences the event before the patient with the lower predicted risk [25]. Values range from 0.5 (random prediction) to 1.0 (perfect discrimination), with higher values indicating better model performance. The C-index is calculated as:

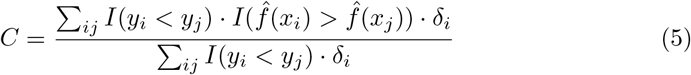

Where *y*_*i*_ is the observed time for patient *i*, 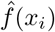 is the predicted risk, *δ*_*i*_ is the event indicator, and *I*() is the indicator function.

#### Integrated Brier Score (IBS)

The IBS assesses both discrimination and calibration by measuring the average squared difference between observed outcomes and predicted probabilities over a range of time points [26, 27]. Lower IBS values indicate better model performance. The IBS is defined as:

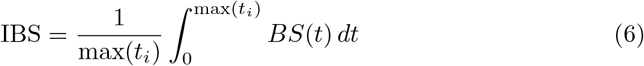

where *BS*(*t*) is the Brier score at time *t*, calculated using inverse probability of censoring weights to account for censored observations.

Mean C-index and IBS values over 10 replicated runs were used to measure predictive accuracy and calibration, providing a robust assessment of model performance.

### Stratification of PIs

PIs were calculated for each patient based on the coefficients of the selected features to quantify individual risk levels. For each survival model, PIs were computed as follows:

#### Cox PH and Elastic Net Cox

The PI is calculated as the linear predictor from the Cox model,

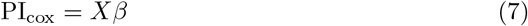

where *X* represents the patient’s covariate values and *β* are the estimated coefficients [28].

#### Random Survival Forest

The PI is derived from the ensemble mortality score,

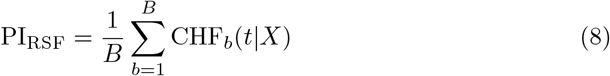

where CHF_*b*_ is the cumulative hazard function estimated from the *b*th tree, and *B* is the total number of trees [22].

#### Survival SVM

The PI is based on the decision function value,

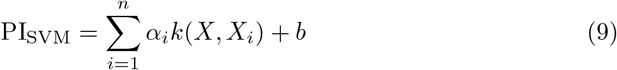

where *α*_*i*_ are the learned weights, *k*(*X, X*_*i*_) is the kernel function, and *b* is the bias term [24].

Risk stratification was performed by dividing patients into risk groups based on PI scores using two different approaches:

#### Two-group stratification

Patients were divided into equal-sized low-risk and highrisk groups (0.5, 0.5) based on the median PI value.

#### Three-group stratification

Patients were divided into low-risk (bottom 1/3), intermediate-risk (middle 1/3), and high-risk (top 1/3) groups based on PI tertiles.

For each stratification approach, Kaplan-Meier survival curves were generated for each risk group, and log-rank tests were performed to evaluate the statistical significance of survival differences between groups [29]. Mean log-rank p-values were computed across the 10-fold cross-validation to summarize model effectiveness in stratifying patients based on survival risk. Additionally, hazard ratios (HRs) with 95% confidence intervals were calculated to quantify the relative risk between groups [30].

### AFT model

To evaluate the prognostic impact of selected gene expression markers, we employed an AFT model with a log-normal distribution, following the framework described by Kalbfleisch and Prentice [31]. Unlike the Cox PH model, which focuses on the hazard function, the AFT model directly models the survival time and assumes that covariates act multiplicatively on survival time, effectively accelerating or decelerating the time to event [32]. Specifically, the model expresses the log of survival time *T* as:

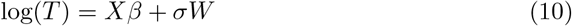

Here, *X* represents the matrix of covariates (i.e., gene expression markers), *β* denotes the regression coefficients, *σ* is a scale parameter, and *W* ∼ *N* (0, 1) is a standard normal error term. The corresponding hazard function for the log-normal AFT model is:

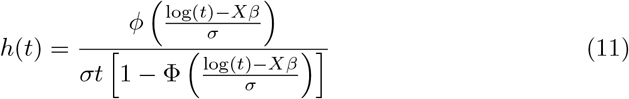

where *ϕ* and Φ are the standard normal probability density and cumulative distribution functions, respectively [33]. HRs were calculated from the model coefficients using the formula:

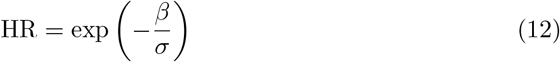

This transformation allows us to interpret each marker’s effect on survival time in terms of relative risk [34, 35]. Specifically, a HR *>* 1 indicates an increased risk (shorter survival time), while a HR *<* 1 indicates a decreased risk (longer survival time).

The log-normal distribution was selected over other parametric options (such as Weibull or log-logistic) based on Akaike Information Criterion (AIC) [36]. We estimated the model parameters using maximum likelihood estimation, with confidence intervals for HRs derived via the delta method [37].

## 3 Results

### 3.1 Performance of markers

To ensure reliability, we selected markers that appeared in at least three out of five cross-validation folds using our random-walk-based approach, resulting in 29 genes. For a fair comparison, we also selected the top 29 genes identified by the regularized feature selection method based on the sum of ranks across all folds. As shown in Figure 1, the netSurvival approach demonstrates comparable performance to other methods in terms of patient risk stratification (measured by IBS) and prediction accuracy (measured by C-index), though it does not outperform them. Notably, our method exhibits consistent performance across all five cross-validation folds.

**Fig. 1.**
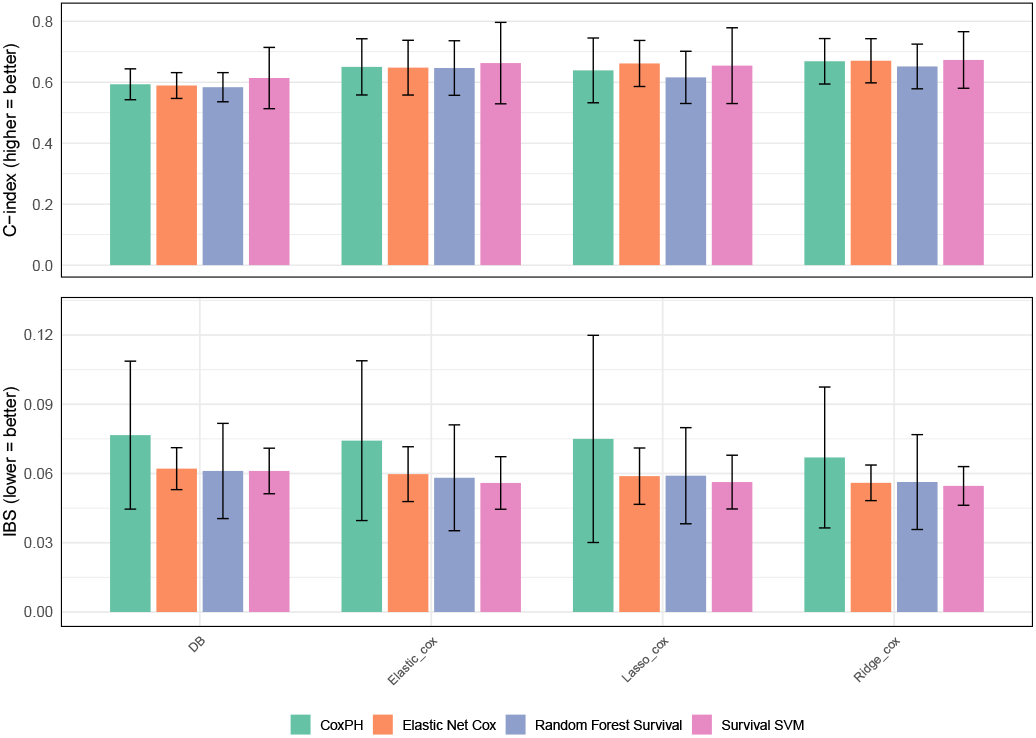
Model performance with different features. Bar plots showing the performance of four survival models: Cox PH, Elastic Net Cox, Random Forest Survival, and Survival Support Vector Machine, across four feature sets derived from different selection methods: Databased-based netSurvival, Elastic_Cox, Lasso_Cox, and Ridge_Cox. The top panel presents the C-index, where higher values indicate better discriminatory power, while the bottom panel displays the IBS, where lower values represent improved overall accuracy. Error bars denote the standard deviation across cross-validation folds.

**Fig. 2.**
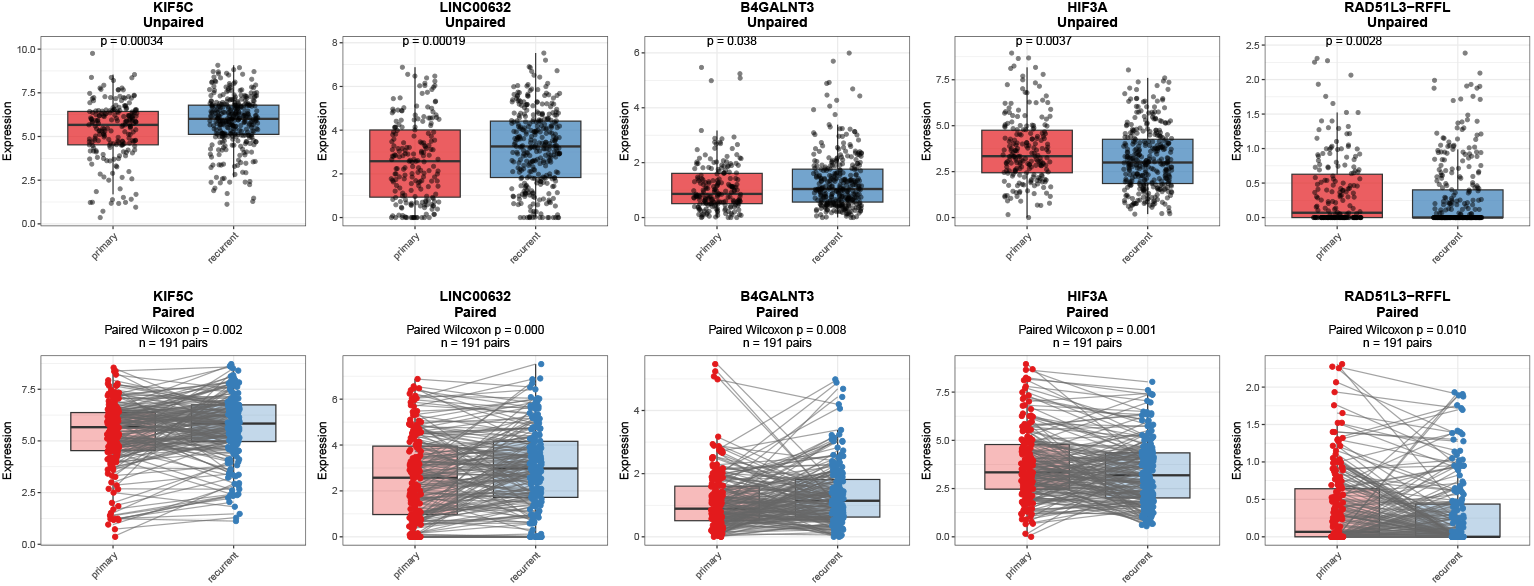
Gene expressions in primary and recurrent samples. Figure shows the expression comparison of five genes (KIF5C, LINC00632, B4GALNT3, HIF3A, and RAD51L3-RFFL) between primary and recurrent IDH wild-type glioma samples using unpaired (top row) and paired (bottom row) statistical tests. Boxplots display gene expression levels with individual sample points overlaid. In the unpaired analysis, a Wilcoxon rank-sum test was used, while the paired analysis applied a Wilcoxon signed-rank test across matched primary–recurrent sample pairs. Significant p-values in both analyses suggest consistent changes in gene expression between primary and recurrent tumors, supporting their potential roles in glioma recurrence.

The table below presents p-values for group comparisons stratified by PIs. While our method’s p-values are not the lowest across all models compared to other approaches, the selected markers remain promising. Although these markers may not be the optimal predictors of patient risk, their competitive performance warrants further exploration.

**Table 1.** Mean and Standard Deviation of Two-Group and Three-Group *p*-values Across Models and Feature Sets.

| Model | Feature Set | Two-Group |  | Three-Group |  |
| --- | --- | --- | --- | --- | --- |
|  |  | Mean | SD | Mean | SD |
| CoxPH | DB | 0.105 | 0.118 | 0.226 | 0.186 |
| CoxPH | Elastic_cox | 0.178 | 0.351 | 0.135 | 0.271 |
| CoxPH | Lasso_cox | 0.133 | 0.260 | 0.138 | 0.207 |
| CoxPH | Ridge_cox | 0.093 | 0.202 | 0.122 | 0.268 |
| Elastic Net Cox | DB | 0.291 | 0.307 | 0.252 | 0.234 |
| Elastic Net Cox | Elastic_cox | 0.112 | 0.187 | 0.222 | 0.374 |
| Elastic Net Cox | Lasso_cox | 0.176 | 0.341 | 0.133 | 0.183 |
| Elastic Net Cox | Ridge_cox | 0.028 | 0.051 | 0.062 | 0.134 |
| Random Forest Survival | DB | 0.712 | 0.346 | 0.275 | 0.257 |
| Random Forest Survival | Elastic_cox | 0.200 | 0.189 | 0.194 | 0.390 |
| Random Forest Survival | Lasso_cox | 0.318 | 0.233 | 0.243 | 0.282 |
| Random Forest Survival | Ridge_cox | 0.211 | 0.249 | 0.055 | 0.086 |
| Survival SVM | DB | 0.254 | 0.275 | 0.176 | 0.257 |
| Survival SVM | Elastic_cox | 0.044 | 0.065 | 0.113 | 0.169 |
| Survival SVM | Lasso_cox | 0.060 | 0.074 | 0.207 | 0.395 |
| Survival SVM | Ridge_cox | 0.201 | 0.279 | 0.172 | 0.378 |

### 3.2 Few-shot prompting

We here check the union set of genes from four different feature selection approaches. The features obtained from all methods can be obtained from Supplementary Table 1. To find out markers with reference support, we implement a few-shot prompting through Grok 3 model. The roles of 15 gene candidates with strong or moderate evidence are listed below.

### 3.3 AFT model evaluation

The AFT lognormal model was employed to assess the association between gene expression levels and survival outcomes, with results presented as maximum likelihood parameter estimates. The model identified several genes with statistically significant effects on survival time (*p <* 0.05), including *FN1, PTK2, CCND2, HIF3A, RAD51L3_RFFL, MAX*, and *EIF4B*. The HRs derived from the model provide insight into the impact of these genes on survival. Specifically, *FN1* (HR = 1.4034, *p* = 0.0183) and *HIF3A* (HR = 1.4907, *p* = 0.0002) were associated with poorer survival outcomes, indicating that higher expression of these genes accelerates time to event, with a 40.34% and 49.07% increased hazard, respectively. Similarly, *EIF4B* (HR = 1.5432, *p* = 0.0013) exhibited a strong adverse effect, with a 54.32% increased hazard for higher expression. Conversely, *PTK2* (HR = 0.7601, *p* = 0.0314), *CCND2* (HR = 0.7781, *p* = 0.0133), *RAD51L3_RFFL* (HR = 0.7928, *p* = 0.0164), and *MAX* (HR = 0.7897, *p* = 0.0418) were associated with improved survival, with HRs below 1 indicating that higher expression of these genes is linked to a slower progression to the event, reducing the hazard by approximately 23.99%, 22.19%, 20.72%, and 21.03%, respectively. These findings highlight the differential prognostic roles of these genes, with *FN1, HIF3A*, and *EIF4B* acting as risk factors, while *PTK2, CCND2, RAD51L3_RFFL*, and *MAX* confer a protective effect on survival.

### 3.4 Correlations with recurrence

Since recurrence is a key contributor to poor prognosis, we examined whether these 67 markers are also associated with tumor recurrence. As shown in Figure 2, the expression levels of five genes, KIF5C, LICNC00632, B4GALNT3, HIF3A, and RAD51L3 RFFL, differ significantly between primary and recurrent groups in both paired and unpaired Wilcoxon tests. Notably, four of these five genes have strong or moderate evidence supporting their functional roles in glioma, as summarized in Table 2, and two show significant HRs in the Cox PH model.

**Table 2.** Summary of genes and their plausible roles in glioma.

| Gene | Evidence | Role in Glioma | Notes |
| --- | --- | --- | --- |
| LINC00632 | Strong | Tumor suppressor, inhibits ALDOA, poor survival with low expression | Direct evidence from TCGA, prognostic marker |
| FN1 | Strong | Promotes invasion, angiogenesis, poor prognosis | Overexpressed, linked to PI3K/AKT pathway |
| PTK2 | Strong | Drives invasion, therapeutic target, associated with Glioma Susceptibility 1 | Well-studied in glioma migration |
| CCND2 | Strong | Cell cycle regulation, proliferation, overexpressed in gliomas | Part of TCGA core pathways |
| LOXL2 | Strong | Enhances invasion, angiogenesis, poor survival | Upregulated in GBM |
| IL13RA2 | Strong | Immune evasion, immunotherapy target, overexpressed in GBM | Prognostic and therapeutic relevance |
| LIF | Strong | Promotes stemness, tumor progression, upregulated in GBM | Linked to cancer stem cells |
| KIF5C | Moderate | Expressed in glioma, potential role in cell division/migration | Inferred from related kinesins, needs more study |
| HIF3A | Moderate | Potential modulator of hypoxia responses, brain-expressed | Less studied than HIF1A/HIF2A, plausible role |
| RAD51L3-RFFL | Moderate | DNA repair and protein turnover, potential therapy resistance | Components suggest relevance, limited direct evidence |
| ATG5 | Moderate | Regulates autophagy, linked to therapy resistance | Critical for cell survival under stress |
| MAX | Moderate | MYC pathway dysregulation, potential proliferation driver | Brain-expressed, needs glioma-specific validation |
| MAPK13 | Moderate | MAPK signaling, regulates proliferation, active in glioma | Part of cancer pathways, plausible role |
| ALDH3A1 | Moderate | Protects cancer stem cells, linked to chemotherapy resistance | Associated with stemness, needs validation |
| EIF4B | Moderate | Regulates translation, supports growth, expressed in cancers | Potential role in glioma proliferation |

**Table 3.**
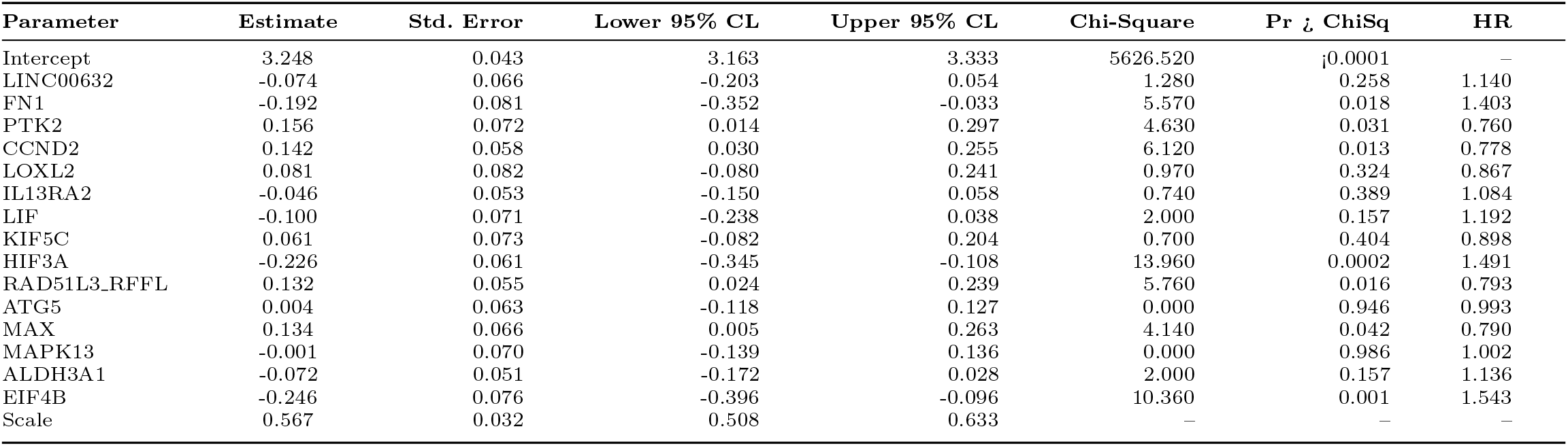
Analysis of Maximum Likelihood Parameter Estimates.

## 4 Discussion

Our comprehensive analysis of gene expression profiles in recurrent IDH wild-type gliomas identified several potential prognostic biomarkers with significant associations with patient survival. By integrating netSurvival with established survival analysis methods, we have uncovered a panel of genes that may contribute to both prognostic stratification and improved biological understanding of IDH wild-type glioma progression.

### 4.1 Interpretation of Key Findings

Our AFT lognormal model revealed that FN1, HIF3A, and EIF4B are significantly associated with poorer survival outcomes, with HRs suggesting substantial increases in risk (40.34%, 49.07%, and 54.32%, respectively). In contrast, PTK2, CCND2, RAD51L3-RFFL, and MAX demonstrated protective effects, with higher expression correlating with improved survival outcomes. These findings align with emerging literature on the biological functions of these genes in glioma pathology.

FN1 (Fibronectin 1) promotes glioma cell invasion and migration by interacting with integrin receptors, particularly *α*5*β*1, contributing to its role as a negative prognostic marker [38]. Similarly, HIF3A (Hypoxia Inducible Factor 3 Alpha) likely contributes to the hypoxic adaptation that characterizes aggressive gliomas, promoting treatment resistance and accelerated disease progression [39]. EIF4B (Eukaryotic Translation Initiation Factor 4B), with the highest hazard ratio among our markers, suggests that dysregulated protein synthesis may be a key driver of aggressive disease behavior in IDH wild-type gliomas.

Conversely, the protective associations observed with PTK2, CCND2, RAD51L3-RFFL, and MAX suggest potential compensatory or tumor-suppressive functions in glioma. PTK2 (Protein Tyrosine Kinase 2) has traditionally been considered prooncogenic in many cancers, but recent evidence suggests context-dependent functions that may explain its association with improved survival in our cohort [40]. CCND2 (Cyclin D2) may reflect a more differentiated cellular state in glioma, consistent with its favorable prognostic association [41].

### 4.2 Recurrence-Associated Biomarkers

The identification of five genes (KIF5C, LINC00632, B4GALNT3, HIF3A, and RAD51L3-RFFL) with differential expression between primary and recurrent tumors provides critical insight into the molecular mechanisms driving recurrence, the key challenge in the management of IDH wild-type glioma. Notably, four of these genes have strong or moderate evidence supporting their functional roles in glioma progression. The dual significance of HIF3A and RAD51L3-RFFL, in both survival outcomes and recurrence patterns, underscores their potential as both prognostic biomarkers and therapeutic targets.

KIF5C promotes glioma progression by enhancing cell proliferation, migration, and invasion through the AKT signaling pathway and cytoskeletal organization [42]. In contrast, B4GALNT3 (Beta-1,4-N-Acetyl-Galactosaminyltransferase 3) may influence invasive properties by glycosylating cell surface proteins [43]. The long non-coding RNA LINC00632 represents an emerging class of regulatory molecules with potential roles in gene expression modulation in glioma [44]. These recurrence-associated biomarkers may provide valuable insights for developing targeted approaches to prevent or delay tumor recurrence, the principal driver of mortality in glioma patients.

### 4.3 Methodological Strengths and Limitations

Our study benefits from several methodological strengths. First, the focus on recurrent IDH wild-type gliomas addresses a critical clinical challenge, as recurrent tumors represent the most therapeutically resistant phase of disease. Second, the integration of multiple feature selection approaches, including netSurvival, provides robust identification of candidate biomarkers. Third, the application of several survival analysis models enhances the reliability of our findings through methodological triangulation.

However, several limitations must be acknowledged. Despite the GLASS consortium providing one of the largest collections of longitudinal glioma data, our sample size remains modest given the molecular heterogeneity of these tumors. Further validation in independent cohorts is essential. Additionally, while gene expression data provides valuable insights, integration with other molecular data types (e.g., methylation profiles, proteomic data) could yield more comprehensive biomarker signatures. Finally, our study does not address potential treatment-induced alterations in gene expression patterns, which may confound the interpretation of recurrence-associated biomarkers.

### 4.4 Clinical Implications and Future Directions

The identified biomarkers hold considerable potential for improving clinical management of IDH wild-type gliomas. In the near term, these markers could enhance prognostic stratification, allowing more precise risk assessment and treatment planning. For example, patients with elevated expression of high-risk markers (FN1, HIF3A, EIF4B) might benefit from more aggressive treatment approaches or closer surveillance. In contrast, while those with favorable expression patterns might be candidates for treatment de-escalation strategies to minimize toxicity.

Beyond prognostication, our findings may inform the development of therapeutic interventions. The negative prognostic association of HIF3A suggests that targeting hypoxia adaptation pathways may be particularly valuable in high-risk patients. Similarly, the protective association of PTK2 raises intriguing questions about its context-dependent functions in glioma that warrant further investigation for potential therapeutic exploitation.

Future research should focus on several key areas: (1) functional validation of these biomarkers through in vitro and in vivo models to establish causal relationships with disease progression; (2) integration with spatial transcriptomic approaches to understand the tumor microenvironmental context of these markers; (3) development of clinically applicable assays for biomarker assessment in routine pathology; and (4) investigation of pharmacological approaches targeting the pathways involving these biomarkers.

## 5 Conclusion

This study identifies a panel of gene expression markers with significant prognostic value in recurrent IDH wild-type gliomas. The differential impacts of these genes on survival outcomes provide insights into the biological heterogeneity underlying clinical behavior in these aggressive tumors. Particularly significant are the biomarkers associated with both survival outcomes and recurrence patterns, which may represent key drivers of disease progression. While further validation is necessary, these findings represent a meaningful step toward improved prognostic stratification and therapeutic targeting in IDH wild-type gliomas, addressing a critical unmet need in neuro-oncology.

## 6 Supplementary information

Supplementary Table 1 of markers selected from four different methods.

## Declarations

### Ethics approval and consent to participate

All patients provided their written, voluntarily informed consent.

### Consent for publication

Not applicable.

### Data availability

The gene expression datasets used in this study are publicly available from the GLASS Consortium: https://www.synapse.org/Synapse:syn17038081/wiki/585622.

### Code availability

All the code can be found through GitHub repository https://github.com/yliu38/netSurvival.

### Author contribution

Y.L., K.K., and J.H. conceived and designed the study. Y.L. performed the computational analysis and wrote the manuscript. K.K., and J.H. edited the manuscript. All the authors reviewed and approved the final manuscript.

## Acknowledgment

None.

## References

[1] Stupp R, et al. Effect of tumor-treating fields plus maintenance temozolomide vs maintenance temozolomide alone on survival in patients with glioblastoma: A randomized clinical trial. JAMA. 2017;318(23):2306–16.

[2] Louis DN, et al. The 2016 World Health Organization Classification of Tumors of the Central Nervous System: a summary. Acta Neuropathol. 2016;131(6):803–20.

[3] Weller M, et al. EANO guidelines on the diagnosis and treatment of diffuse gliomas of adulthood. Nat Rev Clin Oncol. 2021;18(3):170–86.

[4] Ceccarelli M, et al. Molecular profiling reveals biologically discrete subsets and pathways of progression in diffuse glioma. Cell. 2016;164(3):550–63.

[5] Eckel-Passow JE, et al. Glioma groups based on 1p/19q, IDH, and TERT promoter mutations in tumors. N Engl J Med. 2015;372(26):2499–508.

[6] Hegi ME, et al. MGMT gene silencing and benefit from temozolomide in glioblastoma. N Engl J Med. 2005;352(10):997–1003.

[7] Molinaro AM, et al. Genetic and molecular epidemiology of adult diffuse glioma. Nat Rev Neurol. 2019;15(7):405–17.

[8] Neftel C, et al. An integrative model of cellular states, plasticity, and genetics for glioblastoma. Cell. 2019;178(4):835–849.e21.

[9] Ran X, et al. Single-cell transcriptomics reveals the heterogeneity of the immune landscape of IDH–wild-type high-grade gliomas. Cancer Immunol Res. 2024;12(2):232–246.

[10] Park JK, et al. Scale to predict survival after surgery for recurrent glioblastoma multiforme. J Clin Oncol. 2010;28(24):3838–43.

[11] Vredenburgh JJ, et al. Bevacizumab plus irinotecan in recurrent glioblastoma multiforme. J Clin Oncol. 2007;25(30):4722–9.

[12] Wang Q, et al. Tumor evolution of glioma-intrinsic gene expression subtypes associates with immunological changes in the microenvironment. Cancer Cell. 2017;32(1):42–56.e6.

[13] Kim H, et al. Whole-genome and multisector exome sequencing of primary and post-treatment glioblastoma reveals patterns of tumor evolution. Genome Res. 2015;25(3):316–27.

[14] Liu Y, Huse J, Kannan K. Expression graph network framework for biomarker discovery. bioRxiv. 2025. doi: 10.1101/2025.04.28.651033.

[15] Hoerl AE, Kennard RW. Ridge regression: Biased estimation for nonorthogonal problems. Technometrics. 1970;12(1):55–67.

[16] Tibshirani R. The lasso method for variable selection in the Cox model. Stat Med. 1997;16(4):385–95.

[17] Tibshirani R. Regression shrinkage and selection via the lasso. J R Stat Soc Series B Methodol. 1996;58(1):267–88.

[18] Zou H, Hastie T. Regularization and variable selection via the elastic net. J R Stat Soc Series B Stat Methodol. 2005;67(2):301–20.

[19] Simon N, et al. Regularization paths for Cox’s proportional hazards model via coordinate descent. J Stat Softw. 2011;39(5):1–13.

[20] Cox DR. Regression models and life-tables. J R Stat Soc Series B Methodol. 1972;34(2):187–220.

[21] Ishwaran H, Kogalur UB. Consistency of random survival forests. Stat Probab Lett. 2010;80(13-14):1056–64.

[22] Ishwaran H, et al. Random survival forests. Ann Appl Stat. 2008;2(3):841–60.

[23] Van Belle V, et al. Support vector methods for survival analysis: A comparison between ranking and regression approaches. Artif Intell Med. 2011;53(2):107–18.

[24] Pölsterl S, Navab N, Katouzian A. Fast training of support vector machines for survival analysis. In: Appice A, et al, editors. Machine Learning and Knowledge Discovery in Databases. Springer International Publishing; 2015. p. 243–59.

[25] Harrell FE, Lee KL, Mark DB. Multivariable prognostic models: Issues in developing models, evaluating assumptions and adequacy, and measuring and reducing errors. Stat Med. 1996;15(4):361–87.

[26] Graf E, et al. Assessment and comparison of prognostic classification schemes for survival data. Stat Med. 1999;18(17-18):2529–45.

[27] Gerds TA, Schumacher M. Consistent estimation of the expected Brier score in general survival models with right-censored event times. Biom J. 2006;48(6):1029– 40.

[28] Royston P, Altman DG. External validation of a Cox prognostic model: principles and methods. BMC Med Res Methodol. 2013;13:33.

[29] Bland JM, Altman DG. The logrank test. BMJ. 2004;328(7447):1073.

[30] Therneau TM, Grambsch PM. Modeling survival data: Extending the Cox model. Springer; 2000.

[31] Kalbfleisch JD, Prentice RL. The statistical analysis of failure time data. 2nd ed. John Wiley & Sons; 2002.

[32] Klein JP, Moeschberger ML. Survival analysis: Techniques for censored and truncated data. 2nd ed. Springer; 2003.

[33] Collett D. Modelling survival data in medical research. 3rd ed. Chapman and Hall/CRC; 2015.

[34] Wei LJ. The accelerated failure time model: A useful alternative to the Cox regression model in survival analysis. Stat Med. 1992;11(14-15):1871–9.

[35] Carroll KJ. On the use and utility of the Weibull model in the analysis of survival data. Control Clin Trials. 2003;24(6):682–701.

[36] Akaike H. A new look at the statistical model identification. IEEE Trans Automat Contr. 1974;19(6):716–23.

[37] Hosmer DW, Lemeshow S. Applied survival analysis: Regression modeling of time-to-event data. 2nd ed. John Wiley & Sons; 2008.

[38] Serres E, et al. Fibronectin expression in glioblastomas promotes cell cohesion, collective invasion of basement membrane in vitro and orthotopic tumor growth in mice. Oncogene. 2014;33(26):3451–62.

[39] Monteiro AR, et al. The role of hypoxia in glioblastoma invasion. Cells. 2017;6(4):45.

[40] Sulzmaier FJ, Jean C, Schlaepfer DD. FAK in cancer: Mechanistic findings and clinical applications. Nat Rev Cancer. 2014;14(9):598–610.

[41] Koyama-Nasu R, et al. The critical role of cyclin D2 in cell cycle progression and tumorigenicity of glioblastoma stem cells. Oncogene. 2013;32(33):3840–5.

[42] Joyce LJ, Lindsay AJ. A systematic computational analysis of the endosomal recycling pathway in glioblastoma. Biochem Biophys Rep. 2024;38:101700.

[43] Che MI, et al. β1,4-N-acetylgalactosaminyltransferase III modulates cancer stemness through EGFR signaling pathway in colon cancer cells. Oncotarget. 2014;5(11):3673–84.

[44] Wang Y, et al. CRNDE, a long-noncoding RNA, promotes glioma cell growth and invasion through mTOR signaling. Cancer Lett. 2015;367(2):122–8.

